# The phylogenetic signal of extinction through the rise and fall of early vertebrates: field of bullets or clustered strike?

**DOI:** 10.1101/2025.10.27.684355

**Authors:** Joseph Flannery-Sutherland, Amy Tims, Laura Soul, John Clarke, Matt Friedman, Sam Giles

## Abstract

We investigate patterns of phylogenetic selectivity of extinction in early aquatic vertebrates from the Silurian to the Carboniferous, an interval punctuated by one of the ‘Big Five’ mass extinctions and marked by many critical anatomical (e.g. jaws, limbs) and ecological (e.g. macrophagy, terrestrialization) vertebrate evolutionary innovations. Using a new >1300 taxon, formally inferred supertree, we show that phylogenetic extinction clustering in early vertebrates varied through time and between major ecomorphological divisions of the tree. At the end of the Silurian and into the Early Devonian, jawless fishes became marginalized components of vertebrate faunas and show more strongly clustered extinction than jawed fishes, which became ecologically dominant during this interval. Clustered extinction (contemporaneous extinction of close relatives) is typical of most stages of the Devonian, particularly during the Late Devonian extinctions. By contrast, the subsequent early Carboniferous interval of putative recovery is characterized by overdispersed extinction (contemporaneous extinction of distant relatives), consistent with widespread persistence of phylogenetically distinct lineages. This work shows how varying patterns of extinction selectivity pruned the vertebrate tree at the time when the first-order patterns of diversity apparent in modern aquatic vertebrate faunas were first established.

## Introduction

The nearly 200-million-year interval between the end of the Cambrian and the beginning of the Permian records a series of critical evolutionary events in the origin of modern vertebrate diversity: the initial proliferation of armoured jawless fishes (‘ostracoderms’) during the Ordovician (Sansom et al., 2001), the definitive appearance of jawed vertebrates (gnathostomes) during the Silurian (Andreev et al., 2022; Zhu et al., 2022), the origin of terrestrial vertebrates (tetrapods) during the Devonian (Clack, 2012), and major taxonomic restriction of vertebrate faunas associated with the end-Devonian extinction (Sallan and Coates, 2010). Despite these critical events, less quantitative effort has focused on examining patterns and drivers of vertebrate turnover during the mid-Palaeozoic (but see Sallan and Coates, 2010; Friedman and Sallan, 2012; Sallan et al., 2018; Henderson et al. 2022).

While studies of extinction in the fossil record have typically focused on intensity, there is also considerable temporal heterogeneity in patterns of extinction selectivity. Most palaeontological work on selectivity focuses on specific morphological or ecological traits, information which is not easily accessible for many fossil taxa. An alternative approach examines the degree to which extinctions are structured by shared evolutionary history, represented either by taxonomy (Friedman and Sallan, 2012; Hardy et al., 2012; Krug and Patzkowsky, 2015) or phylogeny (Roy et al., 2009; Soul and Friedman, 2017). Although estimates of phylogenetic clustering require data concerning the relationships between taxa, they do not demand detailed trait information. This makes the phylogenetic approach ideal in cases where the taxa of interest are represented by highly fragmentary remains or show such disparate body plans that comparable anatomical traits cannot be measured across all species. This is the situation among mid-Palaeozoic vertebrates, which range from benthic deposit feeders with no paired appendages to species with limbs exploring the margins of continents, and include taxonomically diverse groups known largely from isolated scales, spines, and bones. Interest in key character transformations, however, has resulted in a range of formal cladistic hypotheses for early vertebrates, as well as for major sub-groups (Randle and Sansom, 2017).

Here we leverage this phylogenetic resource to investigate patterns of extinction clustering during the Silurian-Carboniferous history of vertebrates. Because of the disparate habitats and potentially contrasting macroevolutionary pressures affecting early tetrapod evolution, we restrict our study to primitively aquatic vertebrates. We seek to document how phylogenetic selectivity varied between background conditions and intervals of mass extinction and subsequent recovery, alongside whether patterns of clustering differed between jawless and jawed fishes during the establishment of a gnathostome-dominated vertebrate fauna in the Early Devonian (Anderson et al., 2011). Together, these episodes played a critical role in structuring the broad patterns of species richness apparent in extant vertebrates: a diverse gnathostome radiation composed of bony fishes and chondrichthyans, joined by a depauperate assemblage of ecologically and environmentally marginalized jawless fishes (Sallan et al., 2012). Based on expectations from previous literature demonstrating a possible link between phylogenetic selectivity and the ecological consequences of periods of elevated extinction (Droser et al., 2000; Roy et al., 2009; Krug and Patzkowsky, 2015), we specifically test whether: (1) intervals of mass extinction or taxonomic decline are associated with elevated phylogenetic clustering of extinction; and (2) intervals of recovery or evolutionary radiation are associated with phylogenetically random extinction.

## Materials and Methods

### Taxonomic selection

Defining a character set that effectively captures the complexity of anatomical and physiological innovations through early vertebrate evolution is challenging (Donoghue and Keating, 2014), while scoring a sufficiently taxonomically broad sample of early vertebrate taxa would be prohibitively time-consuming. We sampled 127 published character matrices to generate a formal supertree with the requisite taxonomic scope to characterise extinction clustering through the early evolutionary history of aquatic vertebrates.

Matrices were selected to maximise taxonomic coverage of early vertebrate diversity from Stage 2 of the Cambrian to the Serpukhovian stage of the Carboniferous, spanning jawless anaspids, crown cyclostomes, pteraspidomorphs, thelodonts, galeaspids and osteostracans, and jawed placoderms and crown gnathostomes (chondrichthyans, actinopterygians, dipnomorphs, coelacanths and tetrapodomorphs). Following current consensus, we treated conodonts (Euconodonta + Paraconodonta) as stem-group cyclostomes (Goudemand et al., 2011; Murdock et al., 2014; Miyashita et al., 2018; Dearden et al., 2023) and retained any representative operational taxonomic units (OTUs) in our sampled character matrices. As taxonomically broad conodont phylogenies are generally constructed at a suprageneric level, however (e.g., Donoghue et al., 2010), we did not include any character matrices for conodont species themselves. The oldest matrix in our set was published in 1996 but the majority are more recent, with a median publication year of 2015, and we ensured the most recently published matrices (as of 2025) were included for all key clades. In total, 1198 fossil vertebrates are represented across the matrix set, plus five Cambrian-aged early chordate outgroups.

Next, we checked the taxa in our matrix set for corresponding entries in the Paleobiology Database (PBDB). 1677 new occurrences were subsequently added, comprising 321 and 827 previously unrepresented genera and species respectively. Occurrences were added to ensure that the stratigraphic distributions of taxa in the literature were reflected in the PBDB where possible, but we acknowledge that our databasing effort is not spatiotemporally exhaustive. In other cases, genera and species present in the PBDB required revision to new genus or species names, assignment to parent taxa within Chordata, or assignment of their parent taxa to Chordata itself to become available for download. We also attempted to standardise and fix inconsistencies in higher vertebrate taxonomy in the database more generally, an important step as many of the previous taxonomic authorities for early vertebrates were substantially outdated (e.g., the Sepkoski Compendium, 2002).

### Super matrix construction

Older matrices were copied or manually transcribed from paper appendices while more recent matrices were typically available as files in electronic paper supplements or Graeme Lloyd’s archive (graemetlloyd.com/matr.html). All raw matrix files were pre-processed in R (v4.4.1; R Core Team, 2024) to identify and correct any typographical errors in OTU names and character entries, add or correct character ordering information, update or remove junior synonyms and invalid names, and replace tips at genus level or higher with a representative species. Replacements for higher taxa were arbitrarily chosen from suitable species present in other matrices, aside for conodonts where we selected their oldest described species (*Tarimspira zhejiangensis*, Paraconodonta, Cambrian Stage 2 of China) to ensure correct downstream estimation of the node age for vertebrates. Matrices were then re-analysed under equal-weights parsimony using functions from the `TreeSearch` R package (v1.6.0; Smith, 2023) which uses MorphyLib (Brazeau et al., 2017) to handle inapplicable data (Brazeau et al., 2019), and which generally outperforms the alternative method for inapplicable data available in TNT (Simões et al., 2023). Ordered characters were decomposed to a series of binary characters (mathematically equivalent under equal-weights parsimony) then 10 parallel searches initiated, with each search comprising 100 ratchet iterations and 10 tree bisections and reconnections per iteration.

The set of resulting most parsimonious trees (MPTs) from a given analysis was converted to a single maximum parsimony matrix representation (MRP) using functions from the `Claddis` R package (v0.7.0; Lloyd, 2016). Each MRP uses Baum and Ragan (2004) coding to represent all tree bipartitions in the MPT set as ordered characters weighted according to their prevalence in that set, with OTUs scored by presence or absence in each bipartition. These matrices summarise the total uncertainty in early vertebrate relationships recovered during our maximum parsimony analyses and form the basis for constructing a formal super tree.

XML files were generated for each corresponding MRP matrix in R, linking each OTU to a PBDB taxon number, or flagging it for removal in the case of extant OTUs lacking PBDB entries. The complementary sets of XML and MRP files were then supplied to the `Metatree()` function of the `metatree` R package (v0.2.0; Lloyd, 2015) with default settings, which implements the MRP super matrix construction pipeline of Lloyd et al. (2016). Briefly, this pipeline excludes older matrices which do not contribute any unique taxa and weights the contributions of individual MRPs to the final super matrix based on their individual character weights, publication year, degree of data dependency on ‘parent’ matrices in the same MRP set, and degree of conflict between taxonomy and phylogeny within that matrix. This improves the objectivity of super matrix construction (Lloyd et al., 2016), hence why we did not implement any potentially subjective filtering criteria during initial character matrix selection.

### Supertree inference

The MRP supermatrix was subjected to a standard maximum parsimony analysis in TNT (v1.6; Goloboff and Morales, 2023) rather than `TreeSearch` due to the size of the matrix and freedom from inapplicable characters, with search depth `xmult = 10` and 50,000 trees held in memory. A constraint tree (Fig. S1) was used during analysis to enforce the following topology: non-vertebrate chordates as outgroups to the rest of the tree; conodonts and anaspids as sister groups to crown cyclostomes; pteraspidomorphs, thelodonts, galeaspids, osteostracans and pituriaspids as sister groups to jawed vertebrates; arthrodires and antiarchs as sister groups to crown gnathostomes; and chondrichthyans, actinopterygians, coelacanths and dipnomorphs as successive outgroups to tetrapodomorphs. In addition, all genera were constrained to be monophyletic, aside from seven which were explicitly recovered as paraphyletic or polyphyletic in our source matrices (*Groenlandaspis, Onychodus, Psammosteus, Eusthenopteron, Eusthenodon, Traquairaspis, Lampraspis*). A strict consensus tree was then generated from the MPTs recovered under maximum parsimony and any OTUs younger than the Serpukhovian discarded (Fig. S2A).

To maximise taxonomic coverage of early vertebrates, any species present in the PBDB that belonged to genera present in the super tree were grafted in as sister groups (Fig. S2B), aside for instances where genera in the super tree already comprised three or more fully resolved species. While post-hoc grafting incurs the trade-off of decreased phylogenetic resolution in some genera, many others only contain one other species and so complete phylogenetic resolution is maintained in such cases. Grafting was particularly important for thelodonts, as most of their species level diversity is not represented in current character matrices. This decision was also made because inclusion of full species diversity will impact time-scaling by increasing the divergence times required to accommodate the empirical diversity observed within parent genera. In the end, our formal supertree with grafts contains 1345 early vertebrate species, in addition to five chordate outgroups.

### Time-scaling

Previous work has demonstrated that relying on a single time-scaling method may produce spurious results in downstream analyses (Groh et al., 2022). Instead, we time-scaled our supertree using three common *a posteriori* approaches: extended Hedman (EH; Lloyd et al., 2016), three-rate calibration (CAL3; Bapst, 2013), and clockless tip-dating with the fossilised birth-death model (FBD; Zhang et al., 2016). In each case, tip calibrations were derived from the updated PBDB occurrence data for our sample of early vertebrate taxa. All chordate occurrences were downloaded on 16/06/25 via the PBDB application programming interface (Uhen et al., 2023) in R, then subsampled to the species present in our supertree. Collection ages were updated to GTS2020 standard (Gradstein et al., 2020) using the `chrono_scale()` function of the `fossilbrush` R package (v1.0.6; Flannery-Sutherland et al., 2022), then revised using literature searches. PBDB constraints were retained where collection ages could not be updated or refined. Where a collection was assigned to an informal lower (or early), middle, or upper (or late) position within a regional or international chronostratigraphic interval, we took the first to second, second to third, and third to fourth quartiles of the interval duration respectively as the final constraint. Occurrences with stratigraphic uncertainties exceeding 17 million years (the duration of the Emsian, the longest stage in our dataset) were discarded to reduce noise in our downstream analyses, leading to the exclusion of 25 taxa present in our supertree. Lastly, chronostratigraphically suspect entries were identified using the `pacmacro()` function of `fossilbrush` and discarded. Our final occurrence dataset comprises 2267 occurrences representing 1320 species in our super tree, of which 68.7% are singletons, divided into 769 genera across 1176 collections. 1558 occurrences are resolved to stage-level or lower, while a further 1101 occurrences span no more than two stages, with a median collection age uncertainty of 3.35 Ma (median absolute deviation = 1.63 Ma).

EH and CAL3 analyses were implemented in R, using functions from the R packages `strap` (v1.6.1; Bell and Lloyd, 2015) and `paleotree` (v3.4.7; Bapst, 2012), respectively. Each analysis was run 1000 times to generate a sample of time-scaled trees that incorporate stratigraphic uncertainty in occurrence first and last appearance datums (FADs and LADs). During each time-scaling iteration, we constrained the root age (the origin of Chordata) to no older than 575 million years ago in line with estimates from recent total evidence analyses of higher-level metazoan diversification (Carlisle et al. 2024). Collection ages were randomly sampled from within their stratigraphic uncertainty, ensuring that occurrences from a given site received the same random age, following King and Rucklin (2020). Occurrence last appearances were then used for EH, while CAL3 could additionally accept occurrence first appearances.

FBD analyses were implemented in MrBayes (v3.2.7; Ronquist et al., 2012). Input files were generated using the `MrBayesTipDatingNEXUS()` function of `paleotree`. Default function settings were retained as these reflect best methodological practise (Bapst 2012; Matzke and Wright 2016). The youngest tip in the tree (*Cobelodus*, Chondrichthyes, Serpukhovian of Arkansas, USA) was fixed as its youngest possible last appearance age to permit post-hoc transformation of the scaled tree to absolute time. All other tips were split into a pair of terminals, allowing occurrence first and last appearance stratigraphic uncertainties to be assigned as uniform priors on the respective terminals. The root age was assigned a uniform prior of 570 to 580 Ma. To avoid unrealistic drawback of internal nodes due to the long gap between the appearance of conodonts in the Cambrian (the hard minimum for the origin of vertebrates) and phylogenetically widespread emergence of other early vertebrates from the Silurian onwards, we additionally imposed a uniform prior of 470 to 450 Ma on the origin of crown gnathostomes, based on molecular estimates and putative Ordovician chondrichthyan fossils (Sansom et al., 2012; Yu et al., 2024). Markov Chain Monte Carlo (MCMC) was used to estimate the parameters of the FBD model from their posterior distributions. Four replicate analyses were run for each tree for 150 million generations, sampling every 10000, with each replicate analysis comprising four Metropolis-coupled MCMC chains to enable more efficient exploration of parameter space. Model parameters were summarised across all chains. Convergence was identified in all analyses by potential scale reduction factors approaching one (Gelman and Rubin 1992). The first 50% of generations were discarded as burn-in, 1000 random trees selected from the post burn-in posterior sample for analysis, then paired FAD-LAD terminals collapsed back to their original species.

### Extinction clustering analysis

We measured the phylogenetic signal of extinction clustering using two metrics: Fritz and Purvis’ *D* (Fritz and Purvis 2010), and the δ statistic (Borges et al. 2018). Fritz and Purvis’ *D* has been previously used to measure extinction clustering, although its suitability is questionable as the statistic is based on a Brownian threshold model, whereby the evolution of a continuous trait is discretised into a transformation between two binary states (Fritz and Purvis, 2010). While this may be appropriate for some functional, ecological or anatomical traits, extinction versus survival is a strict binary outcome, hence why we also use the δ statistic. This metric is explicitly designed to evaluate categorical trait data based on Shannon entropy provided by ancestral state estimates, under the assumption that a closer association between phylogeny and a categorical trait (i.e., greater phylogenetic signal) decreases the uncertainty in its inferred ancestral states (Borges et al., 2018), although logically the ancestral states of all nodes in this instance must be survival. Extinction clustering analysis was performed in R on the sets of 1000 time-scaled trees from our EH, CAL3 and FBD analyses. The `phylo.d()` function of the `caper` R package (v1.0.3; Orme et al., 2023) was used to measure Fritz and Purvis’ *D*, while we used a custom C++ function and R wrapper to efficiently implement calculation of the δ statistic. Clustering analyses were performed within each stratigraphic stage spanned by a given tree, using all lineages present in each stage, and then for subtrees comprising jawless and jawed taxa.

Finally, we examined several potential correlates of extinction clustering to understand its drivers and biases (Fig. S3). We considered time bin length as a confounding factor affecting the likelihood of a time bin containing taxon LADs, while bin-wise formation and collection counts were taken as proxies for variation in sampling effort through time which may also distort apparent taxon LADs, although we acknowledge that the efficacy of such proxies is limited (Dunhill et al., 2014, 2018). We also calculated the bin-wise mean pairwise great circle distance between collections as a proxy for spatial sampling bias, under the assumption that termination of well-sampled, yet geographically localised sedimentary successions preserving closely related taxa may artificially inflate the intensity of extinction clustering. Our other metrics consider the relationship between diversification dynamics and extinction clustering, namely raw (magnitude) and proportional (intensity) extinction, and mean phylogenetic diversity, calculated bin-wise across each set of 1000 trees from our three time-scaling methods. As each variable was non-normally distributed, we used Spearman’s rank correlation to investigate their relationships with bin-wise mean *D* and δ values, with Benjamini-Hochberg correction for false discovery rates (Benjamini and Hochberg, 1995) due to our large number of comparisons.

## Results

### Supertree structure

Our initial strict consensus supertree is 90.2% resolved (Fig. S2A), although 50,000 MPTs were recovered, indicating that continued analysis would likely recover additional topologies. Tree topology broadly follows early vertebrate interrelationships recovered by previous studies, even though precise ordering of stem gnathostome clades was not enforced during analysis. Pteraspidomorphs are the earliest-diverging total-group gnathostome clade, followed successively by thelodonts, pituriaspids, galeaspids, and osteostracans. Antiarch placoderms, petalichthyids+ptyctodonts, arthrodires, and maxillate placoderms for successive sister groups to the gnathostome crown nodes. While petalichthyid and ptyctodont placoderm monophyly was not enforced during the analysis, they form a clade. ‘Acanthothoracid’ and ‘rhenanid’ monophyly is not supported, and constituent taxa are spread near the base of the placoderm grade. Resolution is poorest amongst thelodonts, with only a few genus-level sister relationships recovered. Polytomies are also prevalent within the highly speciose placoderm genus *Bothriolepis*, as well as between basal chondrichthyan clades. Grafting additional species into genera decreased tree resolution to 76.87%, but resolution amongst deeper nodes in the tree is unaffected (Fig. S2B).

Node ages in the supertree were sensitive to time-scaling method. CAL3 yielded the most conservative root age, estimated as just several million years older than the oldest fossils in the tree (529 – 531 Ma), while EH and FBD yielded root ages in line with our late Ediacaran constraints (580 – 570 Ma). CAL3 also frequently produced branch lengths close to zero in contrast to the other methods, where longer branch lengths and older node age estimates produced smoother phylogenetic diversity (PD) trajectories with greater overall PD throughout the tree (Fig 1). Vertebrate PD increased gradually from the Cambrian to a peak in the latest Silurian, smoothly within EH and FBD trees and stepwise in CAL3 trees (Fig. 1A). PD declined sharply across the Silurian-Devonian boundary, before recovering to a second peak at the end of the Middle Devonian, and declining again throughout the Late Devonian to a low plateau throughout the Tournaisian and Viséan stages of the Mississippian. PD then declined through the remainder of the Mississippian towards the temporal extent of the tree (Fig. 1A). Both jawless and jawed vertebrates contributed to the initial rise in PD, with the former achieving peak diversity by the late Silurian. Both groups also decline across the Silurian-Devonian boundary, but this marked the onset of a terminal decline for jawless groups (Fig. 1B), while the radiation of jawed groups is responsible for the second peak in PD and the trends seen throughout the remainder of our study window (Fig. 1C).

**Fig. 1.**
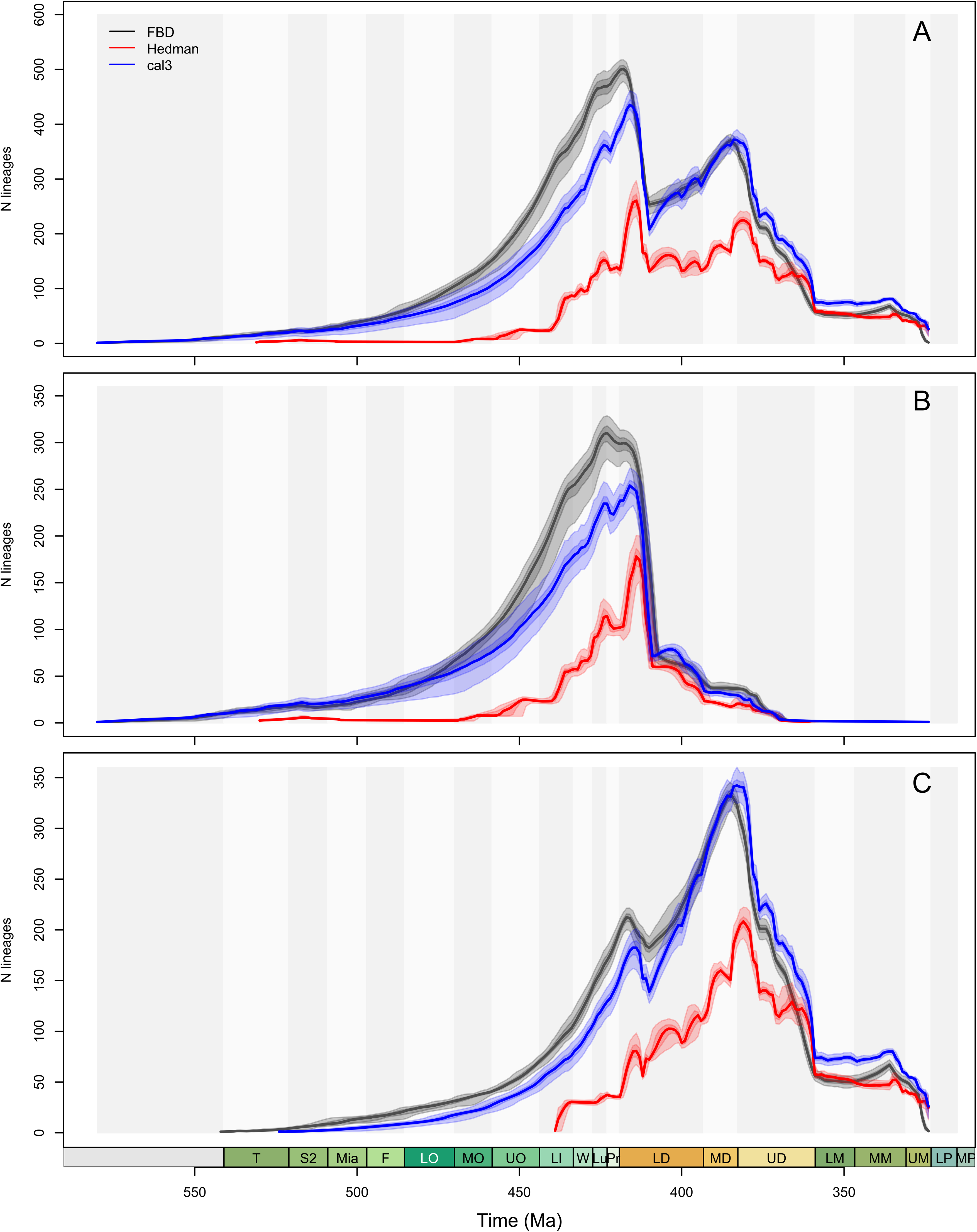
Phylogenetic diversity of early vertebrates. **A.** Lineages through time for all taxa in supertree. **B.** Lineages through time for jawless early vertebrates. **C.** Lineages through time for jawed early vertebrates. Individual lines represent 1000 lineage through time curves from 1000 time-scaled variants of a supertree, with colour groupings indicating the time-scaling method used. Grey divisions denote geological epochs.

### Extinction clustering

Extinction clustering metrics can only be calculated for intervals with a mixture of extinct and surviving lineages. The metrics are also unreliable when the number of lineages available for analysis is small (Fritz and Purvis, 2010; Borges et al., 2018), so we focus here on the time bins where taxon counts generally exceed 50 (Silurian to Mississippian). The strength of extinction clustering varied through time in both jawless and jawed vertebrates and additionally shows sensitivity to tree scaling method, with results from CAL3 trees diverging more strongly from those from FBD and EH trees (Fig. 2, 3; Table S1-9). Conceivably, this relates to the reduced prevalence of ghost lineages in the CAL3 trees due to a high number potentially unrealistic zero-length branches, so we focus here primarily on the results from FBD and EH trees. Conversely, we found no significant relationships between any of our potential correlates of extinction clustering and mean values of *D* and δ (Table S10).

**Fig. 2.**
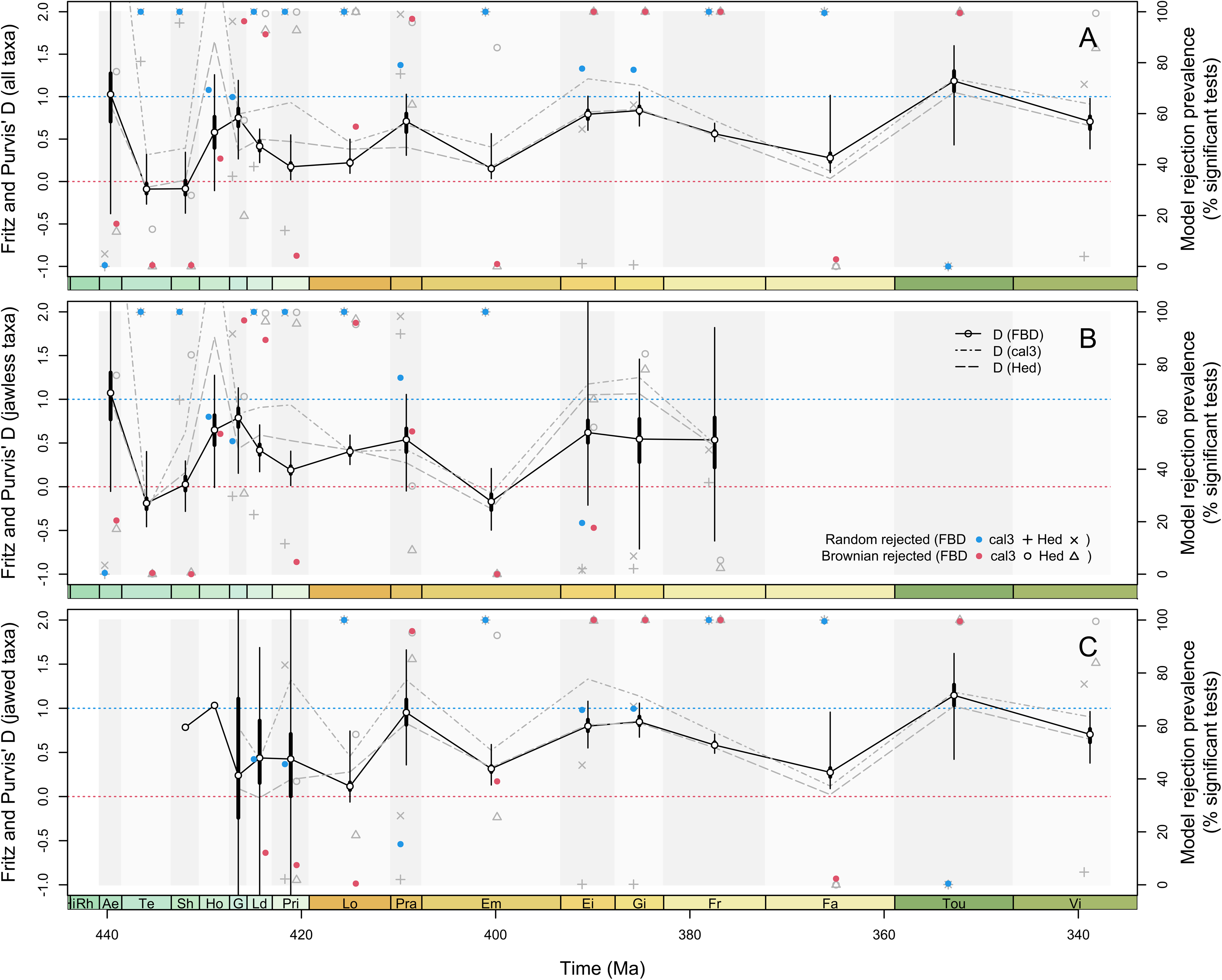
Phylogenetic clustering of extinction in early vertebrates using the D statistic. **A.** Tree-wide extinction clustering. **B.** Extinction clustering for jawless early vertebrates. **C.** Extinction clustering for jawed early vertebrates. Each panel records the stage-wise mean D value across 1000 trees from three different time-scaling methods, in addition to the ranges and interquartile ranges for FBD scaling. Each panel also records the proportion of trees in each stage for which random and Brownian models of extinction clustering were rejected. Grey divisions denote geological stages.

Clustering in jawless taxa in the Aeronian (Silurian) is highly sensitive to branch length, indicated by the wide range of *D* values (Fig 2B). These more closely follow a random structure, but a Brownian model is only rejected across around 20% of the time. Strong extinction clustering is supported in the Telychian, Sheinwoodian and Přidolí (Silurian), while weak clustering is present through the Homerian to Ludfordian (Silurian), with moderate to high rejection of both fully random and fully Brownian models (Fig. 2B). Only the Pragian and Givetian (Devonian) display strong evidence for extinction clustering, with mixed support for solely random or Brownian modes for the remainder of the period (Fig. 2B). The proportion of significant δ values corroborates the strong phylogenetic signal recovered by *D* in the Telychian, Sheinwoodian, Přidolí and Givetian, but additionally suggests prevalent clustering in the Lochkovian (Devonian) and the Ludfordian (Fig. 3B).

**Fig. 3.**
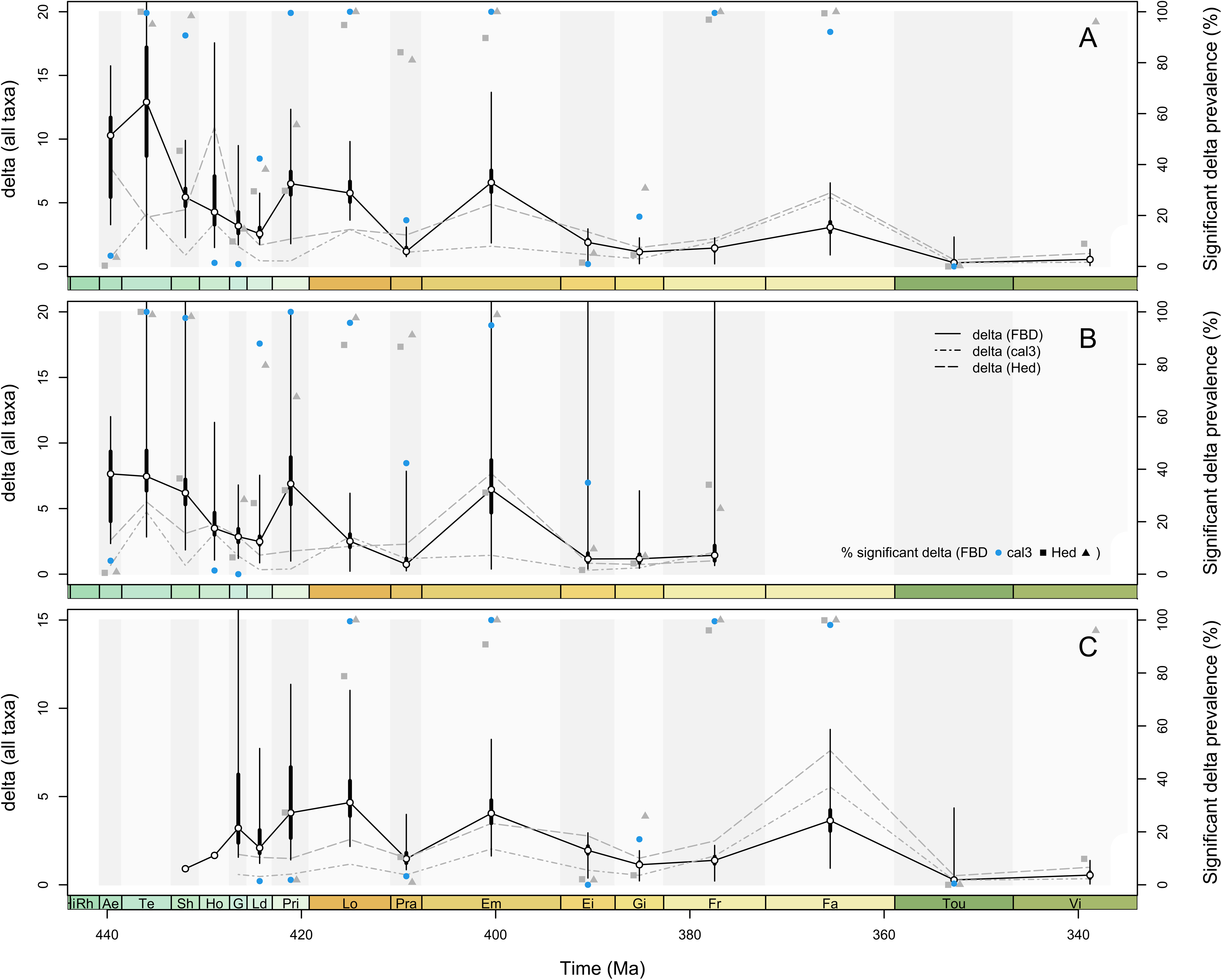
Phylogenetic clustering of extinction in early vertebrates using the. δ **statistic. A.** Tree-wide extinction clustering. **B.** Extinction clustering for jawless early vertebrates. **C.** Extinction clustering for jawed early vertebrates. Each panel records the stage-wise mean δ value across 1000 trees from three different time-scaling methods, in addition to the ranges and interquartile ranges for FBD scaling. Each panel also records the proportion of trees in each stage for which random and Brownian models of extinction clustering were rejected. Grey divisions denote geological stages.

As for jawless taxa, extinction clustering in jawed lineages is supported in the Přidolí (Fig. 2C). Prevalent extinction clustering is also found for the Lochkovian, Givetian and Famennian (Devonian). Conversely, there is strong support for slight phylogenetic overdispersal of extinction in the Tournaisian (Carboniferous). A Brownian model is firmly rejected in the Viséan (Carboniferous), but a random model is only rejected around half of the time, indicating that weak phylogenetic signal may be present during this interval (Fig. 2C). As above, proportions of significant δ values support trends in *D*, indicating extinction clustering in the Lochkovian, Givetian and Famennian, but additionally suggest strong clustering in the Frasnian (Devonian) and Viséan (Fig. 3C).

## Discussion

While we did not identify any diversification-based correlates of extinction clustering, neither did we find that time bin length or geological sampling intensity in either spatial or temporal axes displayed any strong relationship to its phylogenetic signal. As such, we suggest that the trends in extinction clustering reported here are not stratigraphically biased between time bins. Phylogenetic clustering of extinction can provide useful insight into macroevolutionary dynamics in clades through time (Soul and Friedman, 2017). Previous extinction clustering literature has focused on three key questions: whether extinction is random with respect to phylogeny (Hardy et al., 2012; Janevski and Baumiller, 2009; Roy et al., 2009); whether processes acting during times of background extinction are maintained during mass extinctions (Jablonski and Raup, 1995; Krug and Patzkowsky, 2015); and how the degree of phylogenetic clustering observed at a specific extinction events relates to its overall ecological impact (Krug and Patzkowsky, 2015). Our results recover substantial variation in phylogenetic clustering of mid-Palaeozoic early vertebrates through time and between major ecomorphological divisions of the vertebrate tree, providing key insights into all three questions.

### Phylogenetic selectivity of extinction across ecomorphology

The first major pattern to emerge from our analysis is the contrasting degree of extinction clustering between jawless and jawed fishes in the late Silurian and Early Devonian. In each of the three stages of the Přidolí–Pragian interval, our results support that jawless taxa show significantly clustered extinction consistent with a Brownian model. By contrast, there is only evidence for strong extinction clustering among jawed taxa in the Lochkovian, with slight evidence for weak clustering in the Přidolí. This contrast is not readily explained by a difference in taphonomy or preservation rate (a possible source of bias in measurements of extinction clustering; Soul and Friedman, 2017), since jawless and jawed fishes co-occur in many of the same sediments during this interval within our occurrence data, and the presence of large plates of identical mineralogical composition in both groups suggests similar preservation potential.

Significantly, gnathostomes became relatively abundant in the fossil record at this time and a dominant component of vertebrate faunas. By contrast, jawless taxa became less common, with armoured forms represented by substantially reduced diversity largely confined to continental settings with restricted biogeographical distributions (Anderson et al., 2011; Friedman and Sallan, 2012; Sansom et al., 2015; Sallan et al., 2018). Thus, the increased prevalence, elevated morphological diversification, and pronounced ecological innovation of jawed vertebrates during the latest Silurian and earliest Devonian (Anderson et al., 2011; Brazeau and Friedman, 2015; King et al., 2016) was associated with random extinction, while the taxonomic decline and increasing environmental marginalization of jawless fishes was associated with phylogenetically clustered extinction. Whether these divergent patterns are causally linked is unclear. Although the decline of jawless fishes has historically been viewed through the lens of competition with gnathostomes, more recent quantitative efforts have adopted a more circumspect stance (Anderson et al., 2011; Purnell, 2001; Sansom et al., 2015).

### Phylogenetic selectivity of extinction during crisis and recovery

At the scale of the complete phylogeny, the most conspicuous pattern we recover is centred on the Devonian-Carboniferous boundary, corresponding to one of the ‘Big Five’ mass extinctions in Phanerozoic history (Raup and Sepkoski, 1982). The long-recognized Kellwasser and Hangenberg crises of the Late Devonian (McGhee, 1996), particularly the latter, are increasingly interpreted as important episodes in the restructuring of aquatic vertebrate diversity (Sallan and Coates, 2010; Sallan et al., 2011; Friedman and Sallan, 2012; Sallan, 2014; Friedman, 2015; Giles et al., 2017) and triggering of morphological innovation (Sallan and Friedman, 2012; Sallan and Galimberti, 2015). We find strong evidence of significantly clustered extinction during the Late Devonian. This corresponds to the extinction of several species-rich clades, most notably monophyletic radiations within placoderms (e.g. antiarchs, arthrodires) and lobe-finned fishes (e.g. tristichopterids, holoptychiids, some lungfish groups). It has been argued that stronger phylogenetic clustering of extinction leads to greater ecological impact of the event (Krug and Patzkowsky, 2015), and it is certainly true that the pruning of lineages at the end of the Devonian is still apparent in the composition of modern vertebrate diversity.

In contrast to the clustering observed in the Late Devonian, the Carboniferous marks a dramatic shift to non-clustered extinction, beginning with extinctions that are overdispersed relative to a random null model. This indicates a process by which extinction evenly thins a phylogeny, resulting in elevated persistence of distinct evolutionary lineages, rather than coarse pruning of major branches of the tree. This persistence of lineages finds a parallel in patterns reported for taxonomic data, notably the increased longevity of generic cohorts originating in stages following post-Palaeozoic extinctions (Miller and Foote, 2003), and in other phylogenetic studies (e.g. Cretaceous-Palaeogene bivalves; Roy et al 2009). This suggests a degree of conservation of phylogenetic diversity in the recovery phases of extinction events and a ‘winnowing out’ of vulnerable taxa during prior ecologically volatile times.

These patterns, however, are not observed for all mass extinction events. Other phylogenetically explicit investigations have found neither terrestrial vertebrates nor ray-finned fishes show consistent evidence of increased extinction clustering (relative to adjacent pre-or post-extinction intervals) during the end-Permian extinction (Puttick et al., 2017; Soul and Friedman, 2017). Taken together, these results are consistent with an emerging picture of heterogeneity in the relative importance of phylogenetically conserved traits, both during mass extinctions, between different background periods, and between clades (Finnegan et al., 2017; Krug and Patzkowsky, 2015).

### Phylogenetic diversity through early vertebrate evolution

Supertrees have previously been constructed for early vertebrates, but these efforts have focused on representing major clades without attempting to maximise taxonomic coverage of their described diversity, (e.g. 41 taxa in Ferron and Donoghue (2018) or 278 taxa in Yu et al. (2024) with a similar phylogenetic breadth to our analysis). One partial exception to this is the placoderm supertree of Xue et al. (2024), provides complete coverage of described placoderm species diversity by inserting taxa absent from input matrices into higher level polytomies based on taxonomic affiliation. Our tree therefore fills a gap in the literature by providing a phylogenetic hypothesis for most early vertebrate genera that have been included in formal cladistic analyses, alongside their wider species diversity.

While PD may be linked to major biotic events throughout the evolutionary history of a clade (e.g., Condamine et al., 2020), it would be premature to make strong inferences based on our current supertree. Our decision to exclude conodonts is reasonable from a phylogenetic perspective, but means that early vertebrate diversity is systematically underestimated throughout our tree. We estimate that across the Cambrian-Mississippian study interval, our taxonomic coverage of galeaspids, pteraspidomorphs, and osteostracans exceeds 90% at species level, but coverage at genus level is only around 80% for sarcopterygians, 70% for thelodonts, 65% for placoderms, 20% for acanthodians, and 20% for crown chondrichthyans. Similar incompleteness in the PBDB further precludes any comparison with taxonomic diversity in this instance. Despite these limitations, our results can still be considered reflective of extinction selectivity amongst total group gnathostomes rather than vertebrates when conodonts are excluded, provided that incomplete sampling throughout the stem groups and early crown is stratigraphically random. Improving coverage of early vertebrate diversity in the PBDB is therefore key to assessing whether this assumption holds true.

Our divergence time estimates also reveal conflict between time-scaling methods and the ongoing problem of constraining the origin of total-group vertebrates. Estimates from CAL3 diverge most strongly from the other time-scaling methods, which may be attributed to the unreliability of the three rate estimates produced from fossil occurrence datasets containing a high proportion of singletons (Bapst 2012), as is the case for our data and a further reason why we place greater emphasis on the results from our EH and FBD trees.

## Conclusions

Phylogenetic clustering of extinction is variable over time within early aquatic vertebrates, as well as between ecologically distinctive divisions of the vertebrate tree. Jawless fishes show strongly clustered extinction during the latest Silurian and earliest Devonian, the interval during which they became a minor component of vertebrate faunas. By contrast, jawed vertebrates show less clustered or even random extinction during this interval, which appears to represent a time of major taxonomic diversification and ecological innovation in the group. Extinctions were strongly clustered in the final two stages of the Devonian, in advance of the Hangenberg Event. Immediately following this event, during an interval of biological recovery, extinction was significantly overdispersed. These results paint a consistent picture for aquatic vertebrates in the mid-Palaeozoic of reduced extinction clustering during episodes of diversification or recovery and phylogenetically strongly structured extinction during periods of ecological crisis or taxonomic decline.

## Acknowledgements

XXX was funded by Royal Society Dorothy Hodgkin Research Fellowship XXX awarded to XXX who was additionally supported by Royal Society Enhancement Award grant no. XXX. XXX was supported by a Peter Buck Deep-Time Postdoctoral Fellowship. XXX. was supported by the Alexander von Humboldt Foundation and iDiv (funded by German Research Foundation grant XXX), and by European Regional Development Fund grant no. XXX.

## Data and code availability

All code scripts, cladistic matrices and occurrence data used to run our analyses are archived in the paper electronic supplement available at Figshare (10.6084/m9.figshare.30437237). All fossil occurrence data used in this work is also publicly available through the PBDB (paleobiodb.org). TNT is publicly available at lillo.org.ar/phylogeny/tnt/. MrBayes is publicly available at nbisweden.github.io/MrBayes/download.html.

**Fig. S1.**
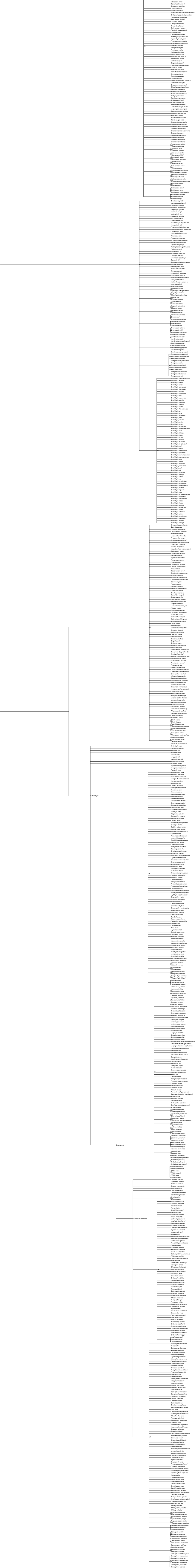
Constraint tree employed during supermatrix analysis.

**Fig. S2.**
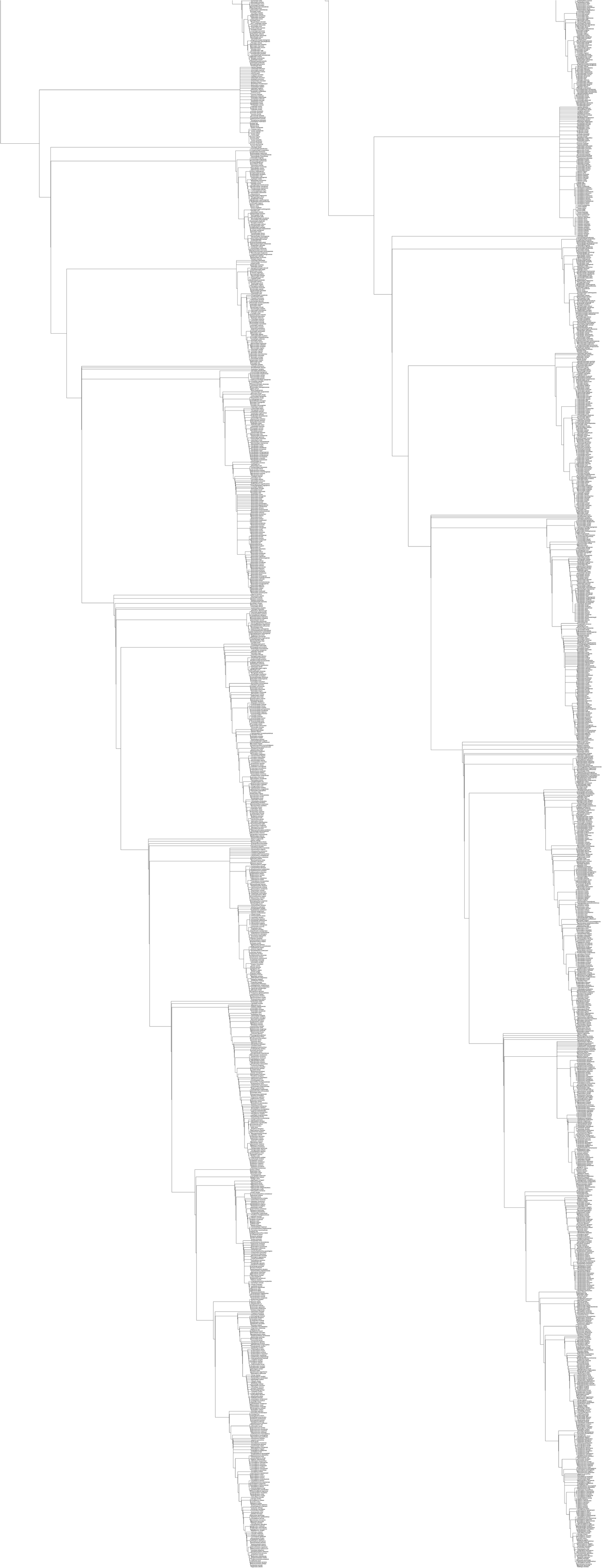
Supertree topology. **A.** Strict consensus from the TNT analysis. **B.** Strict consensus topology with additional species grafted within genus polytomies

**Fig. S3.**
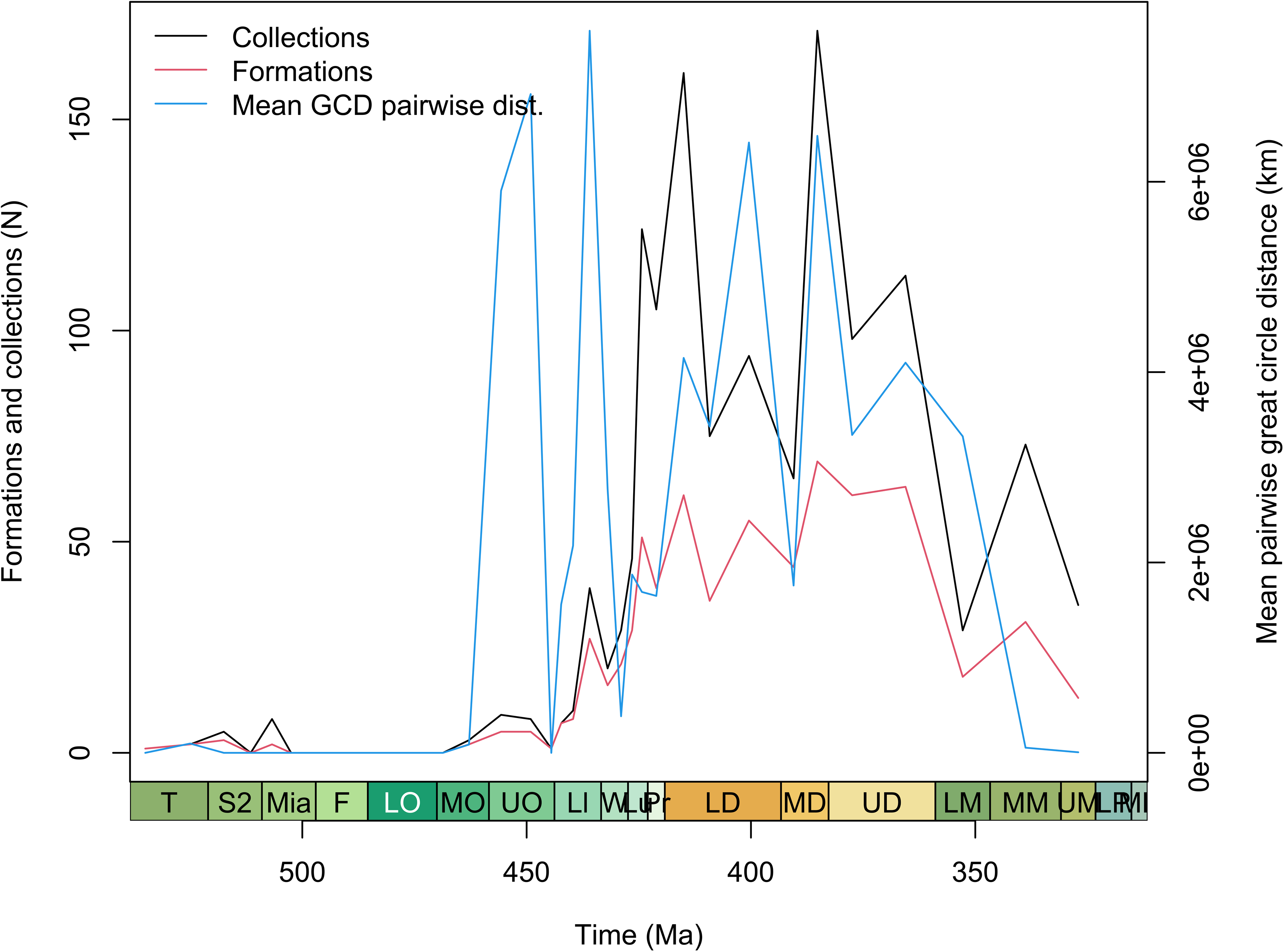
Geological sampling proxies tested for correlations with extinction clustering strength.

## References

1. Andreev, P., Sansom, I., Li, Q., Zhao, W., Wang, J., Wang, C., Peng, L., Jia, L., Qiao, T., Zhu, M. (2022). The oldest gnathostome teeth. Nature, 609, 964–968.

2. Anderson, P., Friedman, M., Brazeau, M., Rayfield, E. (2011). Initial radiation of jaws demonstrated stability despite faunal and environmental change. Nature, 476, 206–209.

3. Bapst, D. (2012). paleotree: an R package for paleontological and phylogenetic analyses of evolution. Methods in Ecology and Evolution, 3, 803–807.

4. Bapst, D. (2013). A stochastic rate-calibrated method for time-scaling phylogenies of fossil taxa. Methods in Ecology and Evolution, 4, 724–733.

5. Bell, M., Lloyd, G. (2015). strap: an R package for plotting phylogenies against stratigraphy and assessing their stratigraphic congruence. Palaeontology, 58, 379–389.

6. Bininda-Emonds, O., Sanderson, M. (2001). Assessment of the accuracy of matrix representation with parsimony analysis supertree construction. Systematic Biology, 50, 565– 579.

7. Brazeau, M., Friedman, M. (2015). The origin and early phylogenetic history of jawed vertebrates. Nature, 520, 490–497.

8. Brazeau, M., Smith, M., Guillerme, T. (2017). MorphyLib: a library for phylogenetic analysis of categorical trait data with inapplicability. Zenodo, doi. 10.5281/zenodo.815371

9. Brazeau, M., Guillerme, T., Smith, M. (2019). An algorithm for morphological phylogenetic analysis with inapplicable data. Systematic Biology, 68, 619–631.

10. Baum, B., Ragan, M. (2004). The MRP method. In: Bininda-Emonds, Olaf, R. P. (ed). Phylogenetic Supertrees: Combining Information to Reveal the Tree of Life. Kluwer Academic, Dordrecht, Netherlands, 17–34.

11. Benjamini, Y., Hochberg, Y. (1995). Controlling the false discovery rate: a practical and powerful approach to multiple testing. Journal of the Royal Statistical Society Series B, 57, 289–300.

12. Borges, R., Machado, J., Gomes, C., Rocha, A., Antunes, A. (2018). Measuring phylogenetic signal between categorical traits and phylogenies. Bioinformatics, 35, 1862– 1869.

13. Carlisle, E., Yin, Z., Pisani, D., Donoghue, P. (2024). Ediacaran origin and Ediacaran-Cambrian diversification of Metazoa. Science Advances, 10, eadp7161

14. Clack, J. (2012). Gaining Ground: The Origin and Evolution of Tetrapods, Bloomington, Indiana University Press, 544p.

15. Condamine, F., Nel, A., Philippe, G., Legendre, F. (2020). Fossil and phylogenetic analyses reveal recurrent periods of diversification and extinction in dictyopteran insects. Cladistics, 36, 394–412.

16. Dearden, R., Lanzetti, A., Giles, S., Johanson, Z., Jones, A., Lautenschlager, S., Randle, R., Sansom, I. (2023). The oldest three-dimensionally preserved vertebrate neurocranium. Nature, 621, 782–787.

17. Donoghue, P., Purnell, M., Aldridge, R., Zhang, S. (2010). The interrelationships of ‘complex’ conodonts (Vertebrata). Journal of Systematic Palaeontology, 6, 119–153.

18. Donoghue, P., Keating, J. (2014). Early vertebrate evolution. Palaeontology, 5, 879–893.

19. Droser, M., Bottjer, D., Sheehan, P., McGhee, G. (2000). Decoupling of taxonomic and ecologic severity of Phanerozoic extinctions. Geology, 28, 675–678.

20. Dunhill, A., Hannisdal, B., Benton, M. (2014). Disentangling rock record bias and common-cause from redundancy in the British fossil record. Nature Communications, 5, 4818.

21. Dunhill, A., Hannisdal, B., Brocklehurst, N., Benton, M. (2018). On formation-based sampling proxies and why they should not be used to correct the fossil record. Palaeontology, 61, 119–132.

22. Flannery-Sutherland, J., Raja, N., Kocsis, A., Kiessling, W. (2022). fossilbrush: An R package for automated detection and resolution of anomalies in palaeontological occurrence data. Methods in Ecology and Evolution, 13, 2404–2418.

23. Finnegan, S., Rasmussen, C., Harper, D. (2017). Identifying the most surprising victims of mass extinction events: an example using Late Ordovician brachiopods. Biology Letters, 13, 20170400.

24. Friedman, M. (2015). The early evolution of ray-finned fishes. Palaeontology, 58, 213– 228.

25. Friedman, M., Sallan, L. (2012). Five hundred million years of extinction and recovery: a Phanerozoic survey of large-scale diversity patterns in fishes. Palaeontology, 55, 707–742.

26. Fritz, S., Purvis, A. (2010). Selectivity in mammalian extinction risk and threat types: a new measure of phylogenetic signal in binary traits. Conservation Biology, 24, 1042–1051.

27. Gelman, A., Rubin, D. (1992). Inference from iterative simulation using multiple sequences. Statistical Science, 7, 457–472.

28. Giles, S., Xu, G., Near, T., Friedman, M. (2017). Early members of a ‘living fossil’ lineage imply later origin of modern ray-finned fishes. Nature, 549, 265–268.

29. Goloboff, P., Morales, M. (2023). TNT version 1.6, with a graphical interface for MacOS and Linux, including new routines in parallel. Cladistics, 39, 144–153.

30. Goudemand, N., Orchard, M., Urdy, S., Bucher, H, Tafforeau, P. (2011). Synchrotron-aided reconstruction of the conodont feeding apparatus and implications for the mouth of the first vertebrates. Proceedings of the National Academy of Sciences of the USA, 108, 8720– 8724.

31. Gradstein, F., Ogg, J., Schmitz, M., Ogg, G. (2020). Geologic Timescale 2020. Elsevier, 1390p.

32. Groh, S., Upchurch, P., Barrett, P., Day, J. (2022). How to date a crocodile: estimation of neosuchian clade ages and a comparison of four time-scaling methods. Palaeontology, 65, e12589.

33. Hardy, C., Fara, E., Laffont, R., Dommergues, J., Meister, C., Neige, P. (2012). Deep-time phylogenetic clustering of extinctions in an evolutionarily dynamic clade. PLoS ONE, 7, e37977.

34. Hedman, M. (2010). Constraints on clade ages from fossil outgroups. Paleobiology, 36, 16–31.

35. Henderson, S., Dunne, E., Giles, S. (2022). Sampling biases obscure the early diversification of the largest living vertebrate group. Proceedings of the Royal Society B, 289, 20220916.

36. Jablonski, D., Raup, D. (1995). Selectivity of end-Cretaceous marine bivalve extinctions. Science, 268, 389–391.

37. Janevski, G., Baumiller, T. (2009). Evidence for extinction selectivity throughout the marine invertebrate fossil record. Paleobiology, 35, 533–564.

38. King, G., Qiao, T., Lee, M., Zhu, M., Long, J. (2016). Bayesian morphological clock methods resurrect placoderm monophyly and reveal rapid early evolution in jawed vertebrates. Systematic Biology, 66, 499–516.

39. King, B., Rücklin, M. (2020). Tip dating with fossil sites and stratigraphic sequences. PeerJ, 8, e9368.

40. Krug, A., Patzkowsky, M. (2015). Phylogenetic clustering of origination and extinction across the Late Ordovician mass extinction. PLoS ONE, 10, e0144354.

41. Lloyd, G. (2015). metatree: Generating Meta-Analytical Phylogenies. R package version 0.1.

42. Lloyd, G. (2016). Estimating morphological diversity and tempo with discrete character-taxon matrices: implementation, challenges, progress, and future directions. Biological Journal of the Linnean Society, 118, 131–151.

43. Lloyd, G., Bapst, D., Friedman, M., Davis, K. (2016). Probabilistic divergence time estimation without branch lengths: dating the origins of dinosaurs, avian flight and crown birds. Biology Letters, 12, 20160609.

44. Matzke, N., Wright, A. (2016). Inferring node dates from tip dates in fossil Canidae: the importance of tree priors. Biology Letters, 12, 1220160328.

45. McGhee, G. (1996). The Late Devonian Mass Extinction: The Frasnian/Fammenian Crisis. New York, Columbia University Press, 307p.

46. Miller, A., Foote, M. (2003), Increased longevities of post-Paleozoic marine genera after mass extinctions: Science, 302, 1030–1032.

47. Miyashita, T., Coates, M., Farrar, R., Larson, P., Manning, P., Wogelius, R., Edwards, N., Anné, J., Bergmann, U., Palmer, A., Currie, P. (2018). Hagfish from the Cretaceous Tethys Sea and a reconciliation of the morphological–molecular conflict in early vertebrate phylogeny. Proceedings of the National Academy of Sciences of the USA, 116, 2146–2151.

48. Murdock, D., Dong, X., Repetski, J., Marone, F., Stampanoni, M., Donoghue, P. (2013). The origin of conodonts and of vertebrate mineralized skeletons. Nature, 502, 546–549.

49. Orme, D., Freckleton, R., Thomas, G., Petzoldt, T., Fritz, S., Isaac, N., Pearse, W. (2023). caper: Comparative Analyses of Phylogenetics and Evolution in R. R package version 1.0.3.

50. Purnell, M., 2001, Scenarios, selection and the ecology of early vertebrates, in Ahlberg, P. E., ed., Early Vertebrate Evolution: Palaeontology, Phylogeny, Genetics and Development: London, Taylor & Francis, p. 187–208.

51. Puttick, M., Kriwet, J., Wen, W., Hu, S., Thomas, G., Benton, M. (2017). Body length of bony fishes was not a selective factor during the biggest mass extinction of all time. Palaeontology, 60, 727–741.

52. Randle, E., Sansom, R. (2017). Exploring phylogenetic relationships of Pteraspidiformes heterostracans (stem-gnathostomes) using continuous and discrete characters. Journal of Systematic Palaeontology, 15, 583–599.

53. R Core Team (2024). R: A Language and Environment for Statistical Computing. R Foundation for Statistical Computing, Vienna, Austria. https://www.R-project.org/.

54. Raup, D., Sepkoski, J. (1982). Mass extinction in the marine fossil record. Science, 215, 1501–1503.

55. Ronquist, F., Teslenko, M., van der Mark, P., Ayres, D., Darlin, A., Hohna, S., Larget, B., Liu, L., Suchard, M., Huelsenbeck, J. (2012). MrBayes 3.2: Efficient Bayesian phylogenetic inference and model choice across a large model space. Systematic Biology, 61, 539–542.

56. Roy, K., Hunt, G., Jablonski, D. (2009). Phylogenetic conservatism of extinctions in marine bivalves. Science, 325, 733–737.

57. Sallan, L. (2014). Major issues in the origin of ray-finned fish (Actinopterygii) biodiversity. Biological Reviews, 89, 950–971.

58. Sallan, L., Coates, M. (2010). End-Devonian extinction and a bottleneck in the early evolution of modern jawed vertebrates. Proceedings of the National Academy of Sciences of the USA, 107, 10131–10135.

59. Sallan, L., Friedman, M. (2012). Heads or tails: staged diversification in vertebrate evolutionary radiations: Proceedings of the Royal Society B, 279, 2025–2032.

60. Sallan, L., Galimberti, A. (2015). Body-size reduction in vertebrates following the end-Devonian mass extinction. Science, 350, 812–815.

61. Sallan, L., Kammer, T., Ausich, W., Cook, L. (2011). Persistent predator-prey dynamics revealed by mass extinction: Proceedings of the National Academy of Sciences of the USA, 108, 8335–8338.

62. Sallan, L., Friedman, M., Sansom, R., Bird, C., Sansom, I. (2018). The nearshore cradle of early vertebrate diversification. Science, 362, 460–464.

63. Sansom, I., Smith, M., Smith, M. (2001). The Ordovician radiation of vertebrates. In: Ahlberg, P. (ed). Major Events in Early Vertebrate Evolution. London, Taylor and Francis, 156–171.

64. Sansom, I., Davies, N., Coates, M., Nicoll, R., Ritchie, A. (2012). Chondrichthyan-like scales from the Middle Ordovician of Australia. Palaeontology, 55, 243–247.

65. Sansom, R., Randle, E., Donoghue, P. (2015). Discriminating signal from noise in the fossil record of early vertebrates reveals cryptic evolutionary history. Proceedings of the Royal Society B, 282, 20142245.

66. Sepkoski, J. (2002). A compendium of fossil marine animal genera. Bulletins of American Paleontology, 363, 1–560.

67. Simões, T., Vernygora, O., de Medeiros, B., Wright, A. (2023). Handling logical character dependency in phylogenetic inference: extensive performance testing of assumptions and solutions using simulated and empirical data. Systematic Biology, 72, 662– 680.

68. Smith (2023). TreeSearch: morphological phylogenetic analysis in R. R Journal, 14, 305–315.

69. Soul, L., Friedman, M. (2017). Bias in phylogenetic measurements of extinction and a case study of end-Permian tetrapods. Palaeontology, 60, 169–185.

70. Uhen, M., Allen, B., Behboudi, N., Clapham, M., Dunne, E., Hendy, A., Holroyd, P., Hopkins, M., Mannion, P., Novack-Gottshall, P., Pimiento, C., Wagner, P. (2023). Paleobiology Database User Guide Version 1.0. PaleoBios, 40, 1–56.

71. Xue, Q., Yu, Y., Pan, Z., Zhu, Y., Zhu, M. (2024). Decline in phylogenetic diversity of Arthrodira (stem-group Gnathostomata) correlates with major Devonian bioevents. Vertebrata Palasiatica, 62, 1–12.

72. Yu, D., Ren, Y., Uesaka, M., Beavan, A., Muffato, M., Shen, J., Li, Y, Sato, I., Wan, W., Clark, J., Keating, J., Carlisle, E., Dearden, R., Giles, S., Randle, E., Sansom, R., Feuda, R., Fleming, J., Sagahara, F., Cummins, C., Patricio, M., Akanni, W., D’Aniello, S., Bertolucci, C., Irie, N., Alev, C., Sheng, G., de Mendoza, A., Maeso, I., Irimia, M., Fromm, B., Peterson, K., Das, S., Hirano, M., Rast, J., Cooper, M., Paps, R., Pisani, D., Kuratani, S., Martin, F., Wang, W., Donoghue, P., Zhang, Y., Pascual-Anaya, J. (2024). Hagfish genome elucidates vertebrate whole-genome duplication events and their evolutionary consequences. Nature Ecology and Evolution, 8, 519–535.

73. Zhang, C., Stadler, T., Klopstein, S., Heath, T., Ronquist, F. (2016). Total-evidence dating under the fossilized birth–death process. Systematic Biology, 65, 228–249.

74. Zhu, Y., Li, Q., Lu, J., Chen, Y., Wang, J., Gai, Z., Zhao, W., Wei, G., Yu, Y., Ahlberg, P., Zhu, M. (2022). The oldest complete jawed vertebrates from the early Silurian of China. Nature, 609, 954–958.

